# Biomonitoring in the Anthropocene: Urban estuary environmental DNA tracks marine fish, terrestrial wildlife, and human diet

**DOI:** 10.1101/2025.09.04.674353

**Authors:** Mark Y. Stoeckle, Jesse H. Ausubel

**Affiliations:** Program for the Human Environment, The Rockefeller University, New York, New York, United States of America

**Author notes:** Corresponding author: (MYS).

## Abstract

Managing human impacts in urban estuaries asks for up-to-date monitoring of marine life. Here we analyze vertebrate eDNA in New York City’s East River, a rocky estuary channel difficult to survey with mechanical gear and subject to wastewater discharge. We collected water samples weekly for one year and applied spike-in metabarcoding to quantify vertebrate eDNA. Replication experiments demonstrated good reproducibility above about 10 eDNA copies/PCR. We propose a fish censusing scale based on absolute eDNA abundance. Local marine fish eDNA followed a classic hollow curve species abundance distribution over four orders of magnitude, with abundant and common taxa comprising about 25% of species and 95% of fish eDNA. There was a 10-fold increase in local marine fish eDNA in summer and seasonal differences among taxa consistent with known phenology. Two fish species were newly abundant in comparison to an eDNA survey at the same site in 2016. Levels of other vertebrate eDNA—domesticated animal, non-fish wildlife, and dietary fish—were correlated with human eDNA levels, consistent with a shared wastewater source. Wastewater eDNA identified the commonest urban mammals, land birds, and household pets. Proportions of dietary animal eDNA in wastewater closely approximated proportions in national consumption statistics, opening a window into human diet assessment. Effort and cost for the weekly shoreline survey were modest. Vertebrate eDNA metabarcoding with spike-in quantification enabled weekly monitoring of urban estuary fish populations, identified overlooked newly abundant species, and reported on terrestrial wildlife and human diet.

## Introduction

Many estuaries are sites of major cities and at the same time essential habitat for wildlife. An urban estuary represents a profound transformation of the physical and living landscape [1,2]. Hazards to the aquatic biosphere may arise from dredging, commercial and residential wastewater discharge, maritime traffic, ballast water disposal, and engineering modifications such as dikes, piers, seawalls, and filling of wetlands [3]. Assessing anthropogenic alterations and mitigations needs up-to-date biological monitoring. Aquatic surveys are constrained by shoreline armoring, navigation restrictions, and submerged structures including cables, tunnels, bridge and pier supports, and rocky reefs.

Environmental DNA may offer a way to advance the science and practice of biomonitoring [4]. The ease and nondestructive nature of collecting water for eDNA raises the prospect of monitoring biodiversity at fine scale in space and time. For instance, there are several metabarcoding primer sets targeting the mitochondrial 12S gene that amplify eDNA from most bony fish species [5–8]. One liter of estuary water typically contains sufficient eDNA to assay at least the more abundant taxa. Hindrances to wider adoption of metabarcoding for fish assessment include persistent uncertainty about how metabarcoding reads relate to eDNA abundance and how eDNA abundance relates to fish abundance [9,10]. Recent reports indicate progress addressing these essential concerns. First, there is increasing evidence that relative metabarcoding reads and eDNA levels are generally proportional to relative fish species abundance [11–15]. Second, this correlation can be improved by weighting metabarcoding reads to adjust for amplification bias, i.e., differences in PCR efficiency among species due to primer mismatch or other factors [16–19]. Alternatively, adding a known quantity of a synthetic DNA template as a standard to each PCR assay (“spike-in”) helps quantify absolute eDNA levels [20–22]. A related approach uses qPCR with metabarcoding primers to quantify total fish eDNA [23]. Quantification offers significant benefits. Absolute eDNA concentrations can be directly compared within and among studies. Spike-in quantification reveals limits to reproducible detection and adjusts for amplification of nontarget DNA [24]. Additional informative work regarding eDNA levels and fish abundance includes mesocosm testing of eDNA shedding and decay, effects of temperature and sunlight exposure on DNA degradation, modeling ocean dispersal including tidal effects, and accounting for allometric scaling [25–29]. Particularly in urban settings, further concerns arise regarding interference from wastewater DNA [30]. eDNA has inherent limitations as a biomonitoring tool, including absent information on life stage, age, health, weight, sex.

In this report we applied spike-in metabarcoding with Riaz 12S gene primers to quantify vertebrate eDNA in New York City’s East River, a tidal strait in the lower Hudson River estuary [31]. The Hudson estuary has a storied history [32,33]. Environmental restoration efforts dating from the Clean Water Act of 1972 have begun to reverse centuries of neglect [34,35]. The East River study site challenges gear-based fish surveys because of the channel’s armored shoreline, irregular rocky bottom topography, and rapid tidal currents [36]. Wastewater permeates the estuary. Like many municipalities, New York City is served by a combined sewer system that conducts household sewage and stormwater from street runoff to underground reservoirs and then to treatment plants [37]. When conduits are overloaded, as frequently occurs in New York City after even modest rainfall, the effluent empties at combined sewer overflow (CSO) outfalls into waterways. CSO outfalls in New York City currently discharge about 18 billion gallons of untreated wastewater into the estuary annually. Although significant, this represents an 80% reduction in CSO discharges since 1985, thanks to more than $40 billion in infrastructure improvements [38]. We test two hypotheses: first, that estuary eDNA quantifies local marine fish populations without interference from wastewater DNA and second, that wastewater DNA usefully reports on other aspects of urban environment.

## Materials and Methods

Replicate 1 L water samples were collected weekly at an East River shoreline site, used in prior studies, beginning May 2, 2024 through May 1, 2025 (Fig 1) [39]. As the shoreline is armored and elevated about 2 meters above water level, paired samples were obtained with separate throws of a bucket on a rope and transferred at site to laboratory bottles. Given tidal current up to 3 m/sec, estimated average water surface distance between samples was about 100 m (range 0 m – 400 m) [36]. Samples were brought to the laboratory within 15 min and each liter was filtered separately using a 4.5 cm, 0.45 μM pore size nitrocellulose membrane (Millipore) and wall suction. Once complete, filters were folded to protect retained material and stored in 15 mL centrifuge tubes at −20°C. Between uses, collection and filtration equipment were washed extensively with tap water and air dried. DNA was extracted from filters within three months using Qiagen DNeasy PowerSoil Pro Kit and eluted in 100 μl of Buffer 6. DNA yield was measured with Qubit and samples were stored at −20°C. Negative field controls consisted of 1 L samples of laboratory tap water filtered and extracted for DNA using the same equipment as for field samples.

**Fig 1.**
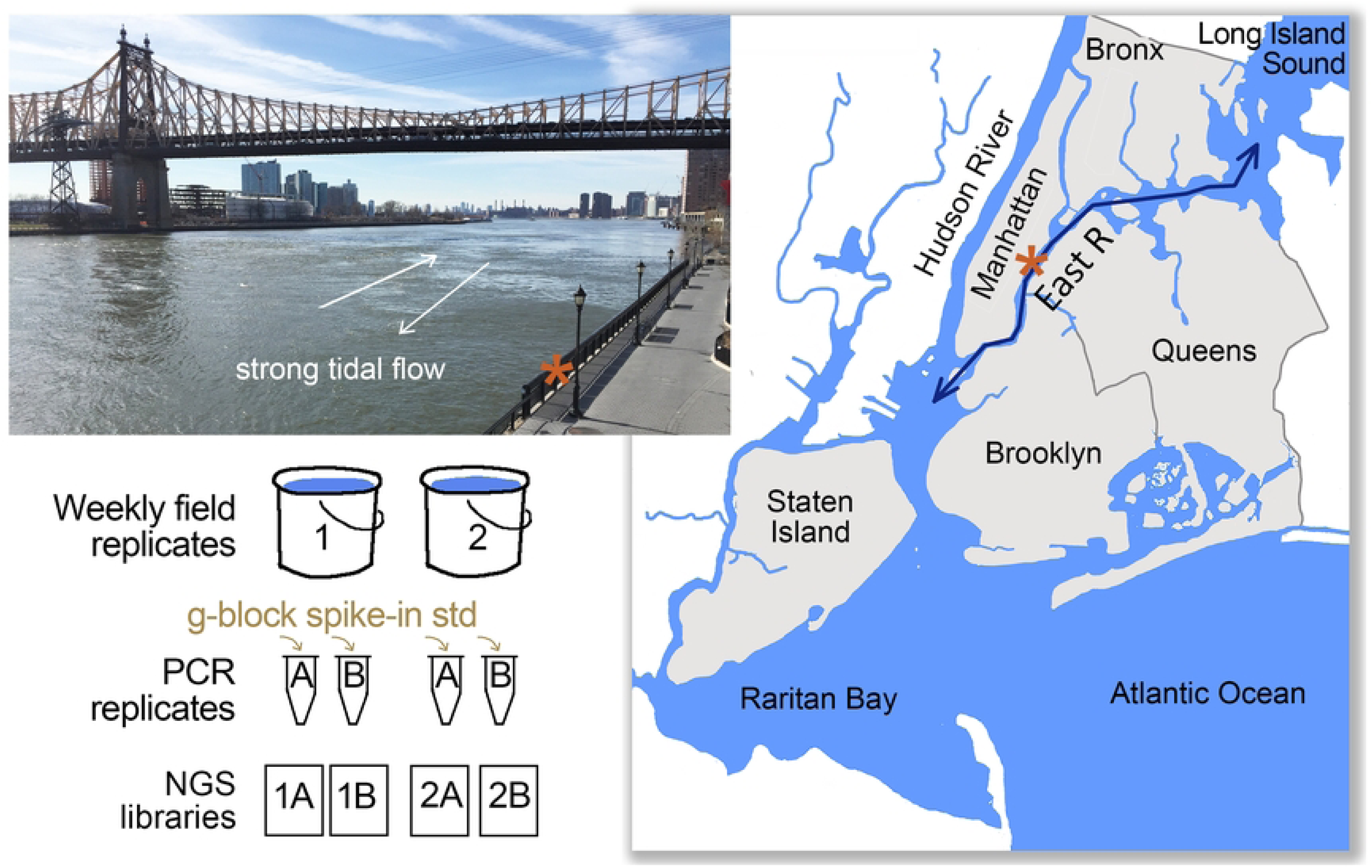
Survey design. Upper left, Collection location in New York City’s East River, a tidal channel between Long Island Sound and New York Harbor. Asterisk marks collection site. Right, map of lower Hudson River estuary and surrounds. Lower left, collection and analysis strategy. Map generated in Photoshop using USGS templates (https://apps.nationalmap.gov/viewer/). Photo credit Mark Stoeckle.

PCR was done as previously described except that the spike-in standard was a 768 bp synthetic gene block (IDT), based on native ostrich 12S DNA amplicon standard in prior study (S1 Fig) [40]. Unlike the native ostrich 12S sequence, the synthetic gene block has M13 tails and three bases modified to match the MiFish-U forward primer [6]. Primary PCR assays were carried out with TaKaRa Titanium Taq hot start High-Yield EcoDry premix in 25 μL volume with 200 μM tailed Riaz primers, 250 copies gene block standard, and 5 μL DNA or 5 μL reagent-grade H2O, the latter as PCR negative control. Unlike other commonly used primer sets for fish metabarcoding, Riaz 12S primers are effective for most vertebrates, as 12S binding sites are conserved not only in bony fish but also in mammals and birds [5]. DNA samples were amplified in replicate and negative field and PCR controls were included in each amplification set. Primer sequences and thermal cycling conditions are in S1 Table. After amplification, 5 μL of reaction mix were run on 2% agarose gel with SyberSafe to assess products and the remaining 20 μl were diluted 1:20 in Buffer EB and stored at −20°C. Following procedure described in prior studies, 5 μL of diluted primary PCR product were indexed with Nextera XT Index Kit v2 Set A and Cytiva PuReTaq Ready-to-Go PCR beads in 25 μl volume. To assess products, 5 μL of index reaction mix were run on a 2% agarose gel, and the remaining 20 μL were pooled, cleaned with AMpure beads at 1:1, and resuspended in Buffer EB. Pooled libraries were sequenced at AZENTA using MiSeq, 2 x 150 bp, 10% PhiX, and 7.5 GB depth. Each library represented a single PCR run on a single DNA sample. The 192 field libraries plus 21 controls (11 field, 10 PCR) were analyzed in four sequencing runs together with unrelated libraries. To assess potential primer bias, a set of DNA samples (64 field samples plus negative controls) were amplified with MiFish-U-F/R2 primers and ostrich g-block spike-in as described above. MiFish-U-F/R2 primers have an extra 3’ base in reverse primer to reduce amplification of bacterial 16S DNA [40]. Primer sequences and thermal cycling parameters are in S1 Table. The protocol was otherwise the same as for Riaz primer amplifications. Pooled libraries were cleaned with AMpure and sequenced at AZENTA on a MiSeq, 2 x 250 bp with 10% PhiX.

Bioinformatics used a DADA2 pipeline as previously described [24]. DADA2 output files were transferred to Excel. Amplicon sequence variants (ASVs) were filtered to exclude detections representing less than 0.1% of total reads for a given ASV. ASVs were identified by 100% match to a local library of reference sequences representing local marine fish, local freshwater fish, nonlocal fish, non-fish wildlife, domesticated animal, and human (S2 Table). Unmatched sequences were manually submitted to GenBank using BLAST; additional matches were added to the reference library. For each library, reads per copy of ostrich standard were calculated. This proportion was then applied to convert reads to eDNA copies for all ASVs in that library (reads per ASV ÷ reads/copy standard = copies per ASV). As previously reported some local marine fish species shared Riaz segment 12S sequences, and some species were represented by more than one ASV (S2 Table). Results are expressed as eDNA copies per PCR assay or as per liter water sample, the latter obtained by multiplying copies per PCR by 20 to account for the proportion of DNA extract used for each PCR. For MiSeq fastq files, the pipeline was adjusted to accommodate the longer amplicon. Statistical tests were performed in GraphPad Prism 10.5.0.

## Results

### Vertebrate eDNA by category

A total of 96 samples were obtained on 48 collecting days. eDNA was successfully extracted and amplified from all water samples and negative controls, generating 192 field sample and 21 negative control NGS libraries. ASVs in DADA2 output files were identified to species and assigned to vertebrate categories as described in Materials and Methods. Sequencing depth appeared sufficient to detect single copy eDNA. Average sequencing depth was 116,983 vertebrate reads per library (range 21,045 – 275,753); average reads per eDNA copy gene block standard were 88 (range 4 – 326) (S3 Table). Local marine fish eDNA was detected on all days (average copies/L, 12,507; range 1,161 – 58,738) but accounted for less than half of total vertebrate eDNA (Fig 2, S6 Table). The majority was human eDNA (average copies/L, 27,629; range, 1,295 – 279,273). Domesticated animal and other categories of vertebrate eDNA were commonly present. Negative control libraries yielded low levels of human (average copies/L, 24; range 0 – 173) and domesticated animal (average copies/L, 15; range 0 – 143) eDNA. (Fig 2, S4 Table). Other categories of vertebrate eDNA were not detected in negative controls.

**Fig 2.**
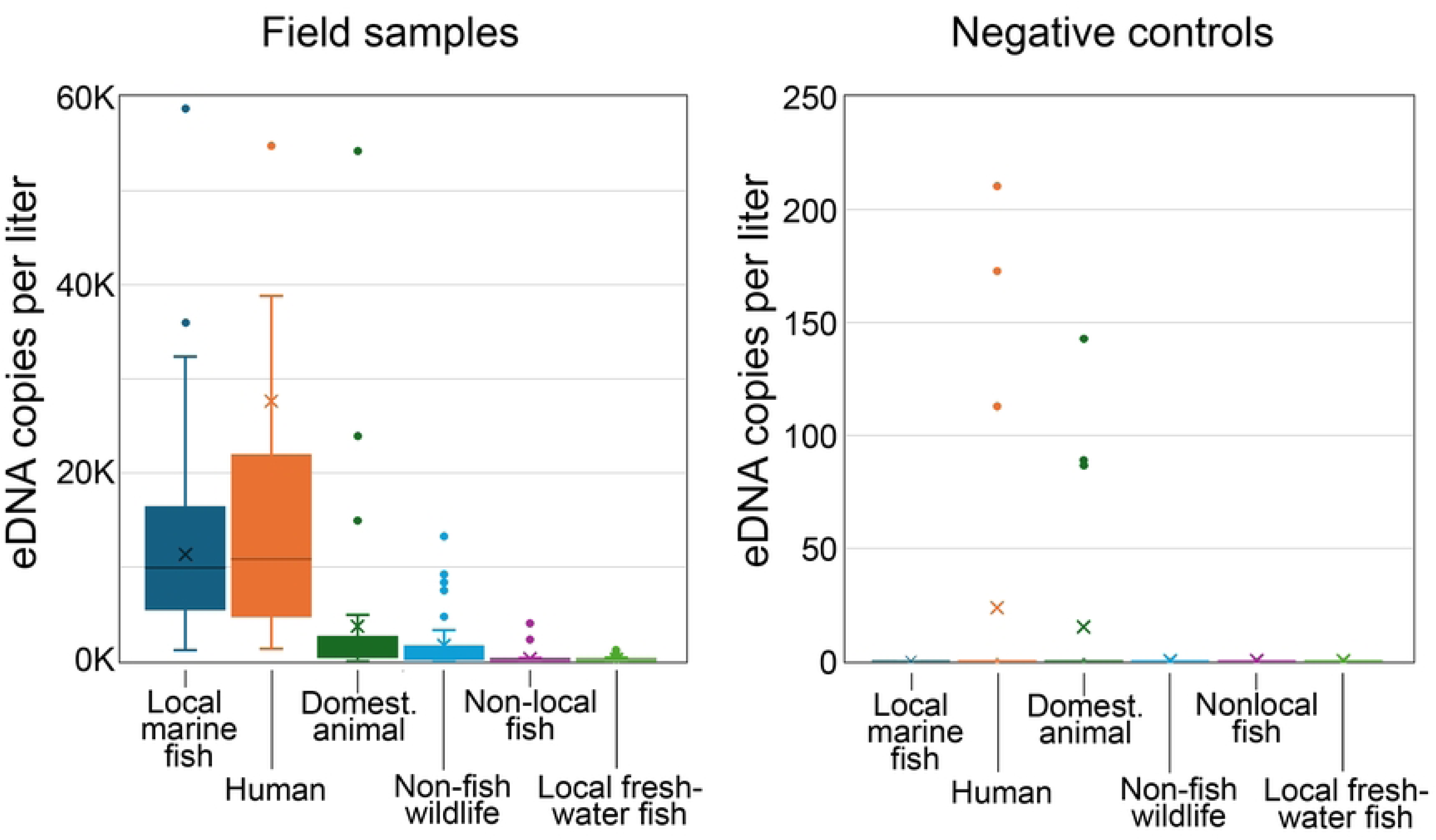
Vertebrate eDNA abundance in field samples and negative controls. Values represent daily averages (left) or individual negative controls (right). Note scale differs between graphs. Box indicates 25th-75th percentile, whiskers, 1.5x interquartile range, and outliers are shown as points; x marks average and line is median. Graph at left does not display outliers in human eDNA; plot otherwise represents values for complete dataset (S4 Table).

### Local marine fish

#### eDNA copy number reproducibility

Species detection and eDNA copy number were largely reproducible in single PCR replicates, particularly for more abundant eDNA (S5 Table, S2 Fig). For species with more than 10 copies/PCR, most (414/443; 94%) were detected in replicate library, whereas for those with fewer than 10 copies/PCR, only about half (124/278; 45%) were present. Excluding nonreplicated detections, the average absolute fold-difference in copies per species was modest (average, 2.0; standard deviation (SD), ± 2.5). As expected, there were larger differences between paired field collections (S5 Table, S2 Fig). About half of field sample detections above and below 10 copies/PCR were present in replicate library (225/443 (51%) and 125/278 (45%), respectively). Excluding non-detections, the average absolute fold-difference in copies per species was 3.1, SD, ± 8.2. Pooled PCR and field replicate datasets were reproducible over a wide concentration range (Fig 3). The average absolute fold-difference in copies per species was 1.3 (SD ± 0.4) and 1.5 (SD ± 1.3), respectively (S6 Table). Amplification with MiFish-U-F/R2 primer set yielded copy numbers per species similar to that with Riaz primers (Fig 3; S7,S8 Tables; S3 Fig). One exception was cunner (*Tautogolabrus adspersus*), which amplified weakly with MiFish-U-F/R2 primers; this discrepancy was predictable based on primer mismatch (S4 Fig).

**Fig 3.**
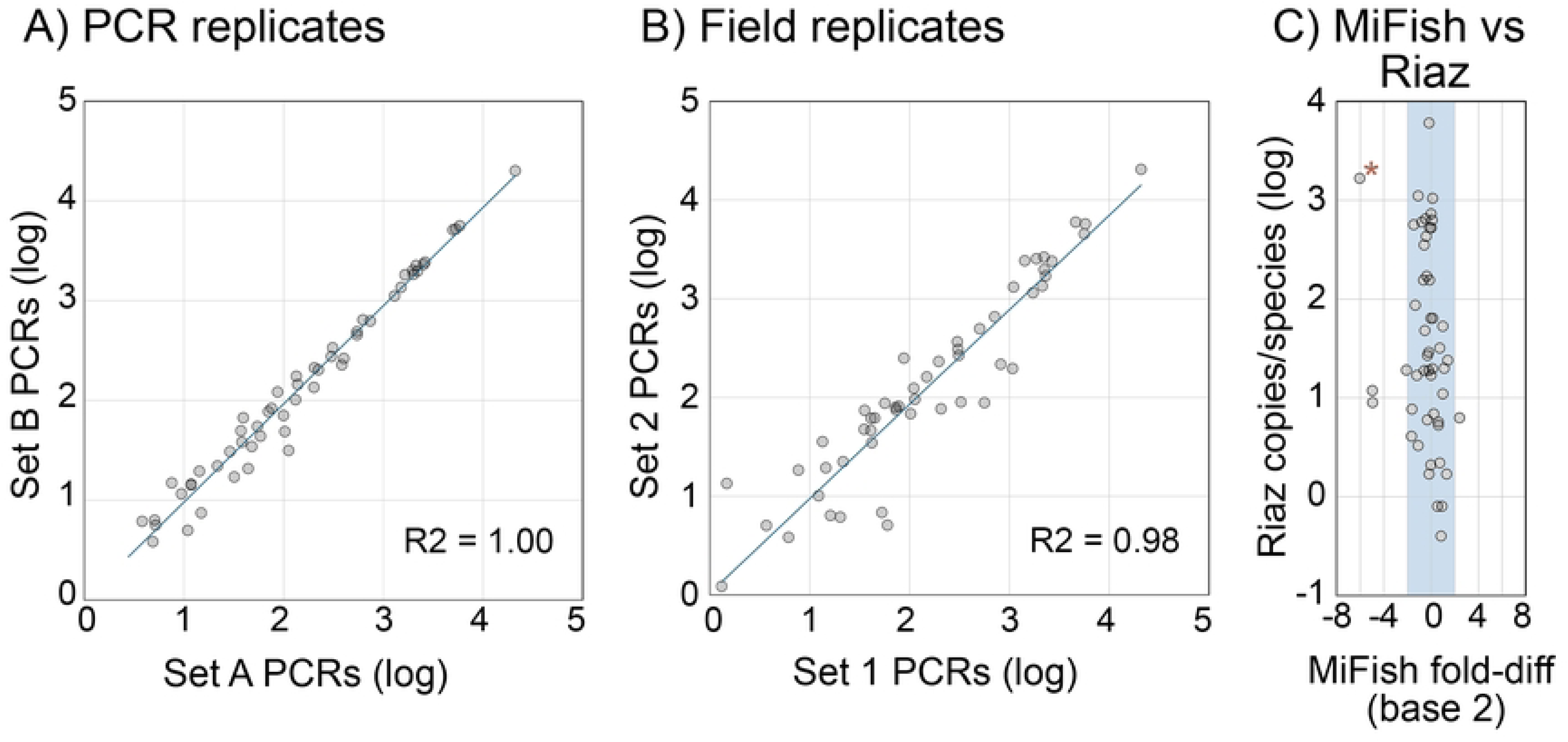
Reproducibility of pooled PCR and field replicates. Each point represents one local marine fish species in pooled sets of A) PCR or B) field replicates (96 PCRs/set). Sets are named as in Fig 1. Values are copies per pool. C) Fold-difference copies/species for pooled datasets generated with MiFish-U-F/R2 as compared to Riaz primers. Shading covers ± 4-fold difference. Asterisk indicates cunner (*T. adspersus*). Data sources in text.

#### Species abundance distribution (SAD)

Seventy-one local marine fish species were detected (S3 Table). Species yearly average eDNA abundance followed a hollow curve distribution ranging over four orders of magnitude (Fig 4). We propose an abundance scale based on order of magnitude differences in absolute eDNA concentration as shown in Fig 4. Abundant species were commonly detected and when detected, present in many copies (S5 Fig). Conversely, rare species were rarely detected and when detected, present in few copies. These two properties generated the hollow curve SAD. Abundant and common species (n = 16; 23% of species) were mostly reef-associated, consistent with the rocky nature of the East River site and accounted for the great majority (95%) of fish eDNA (Table 1).

**Fig 4.**
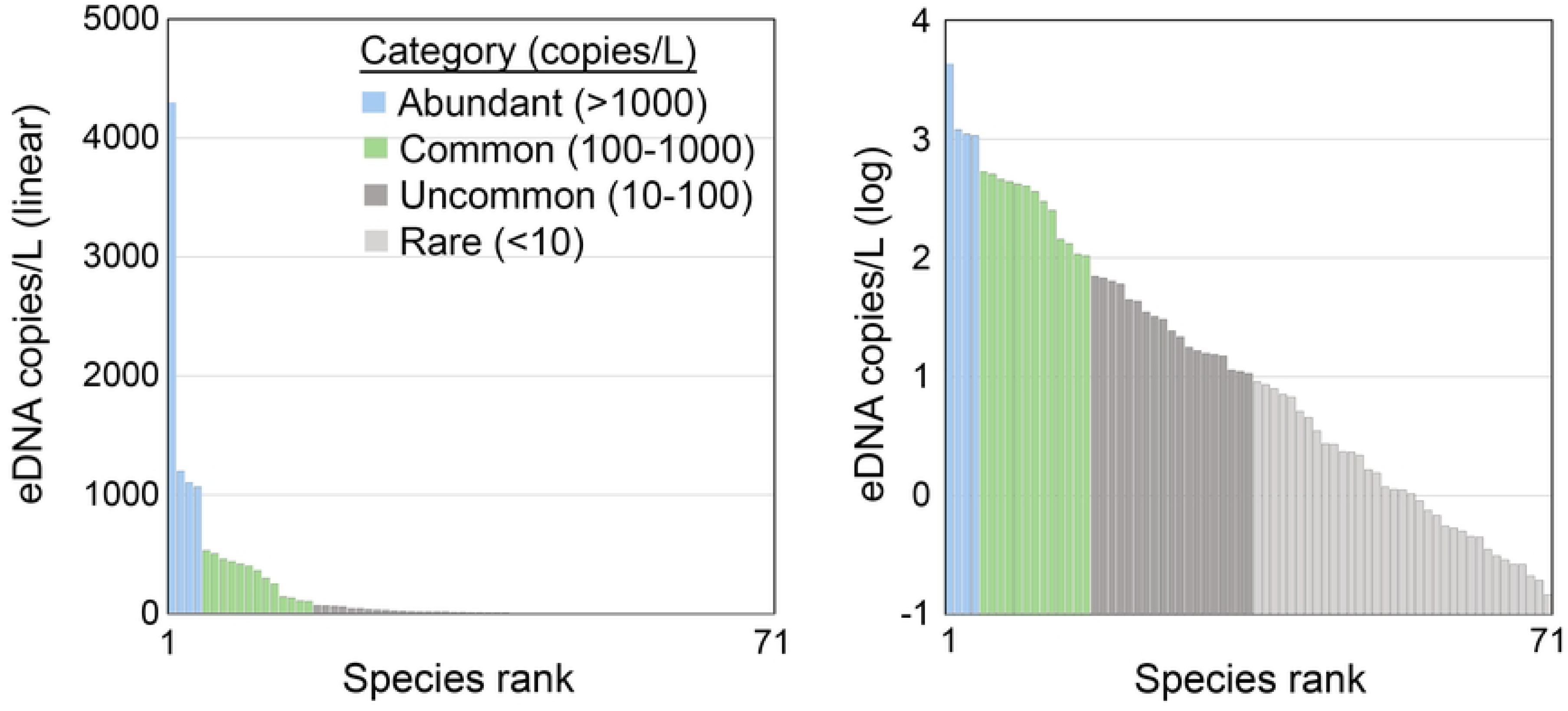
Local marine fish SAD and proposed abundance categories. Each column represents one species. Species are rank ordered by average yearly abundance and shown in linear and log scale.

**Table 1.**
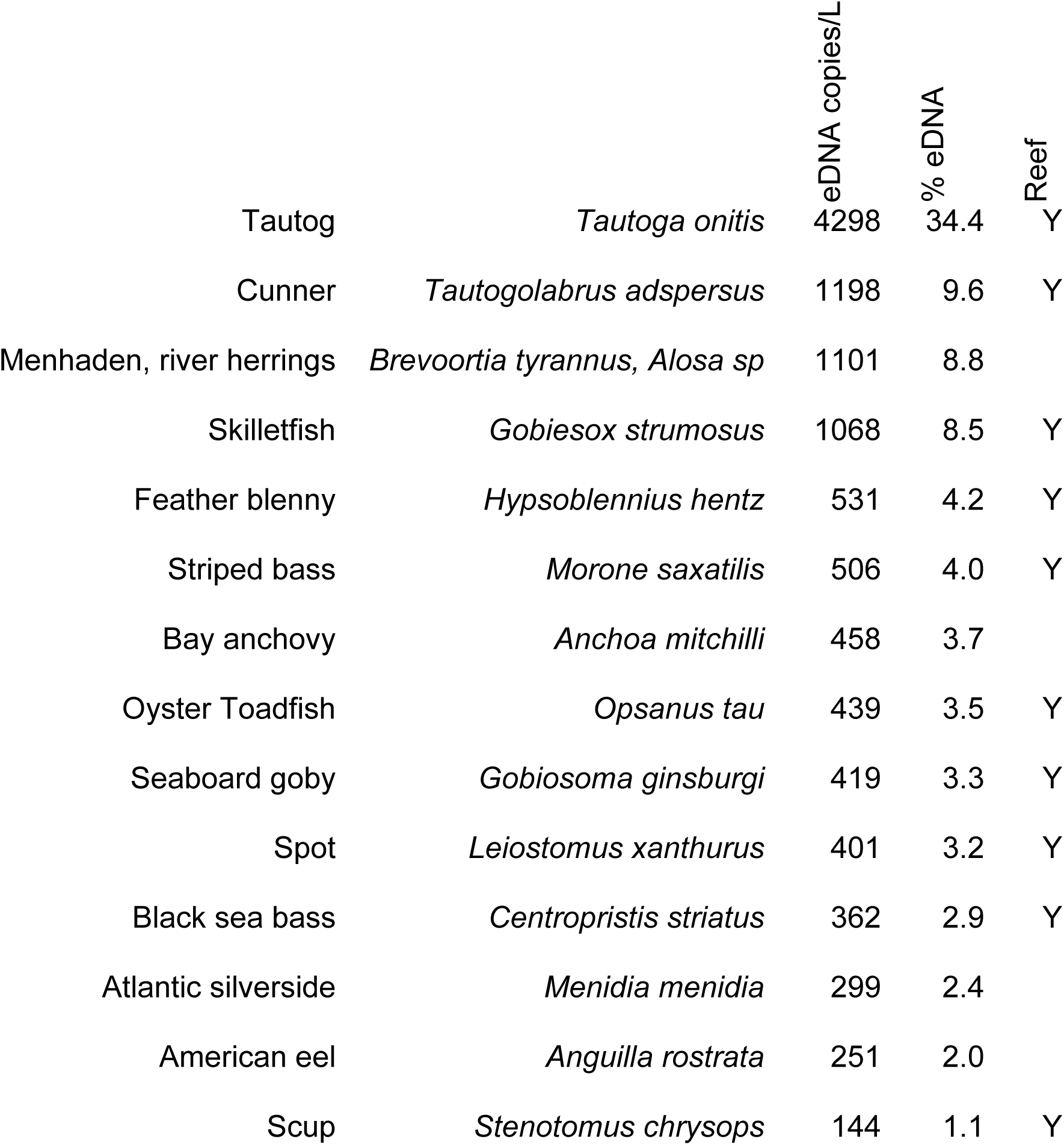
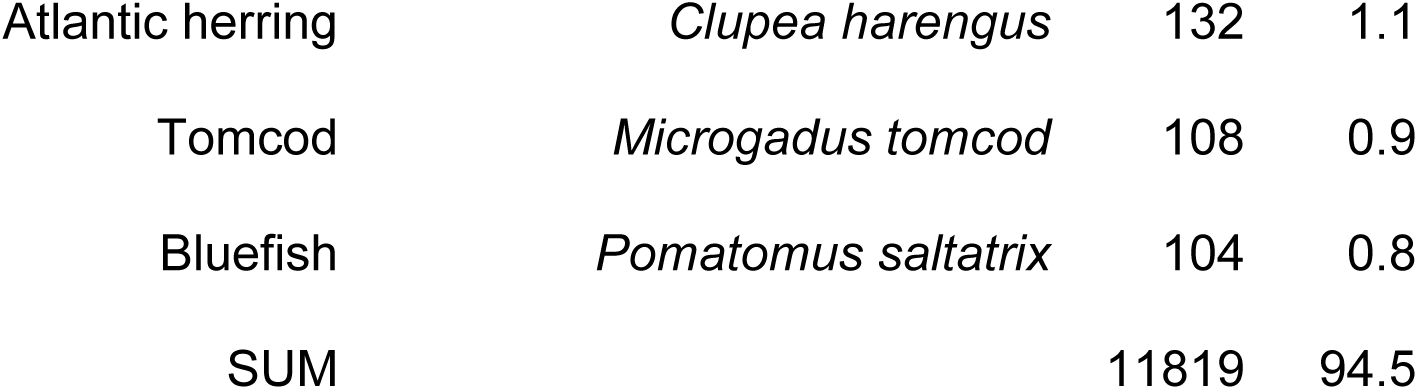
Abundant, common local marine fish. Species are ranked by decreasing abundance.

|  |  | eDNA copies/L | % eDNA | Reef |
| --- | --- | --- | --- | --- |
| Tautog | *Tautoga onitis* | 4298 | 34.4 | Y |
| Cunner | *Tautogolabrus adspersus* | 1198 | 9.6 | Y |
| Menhaden, river herrings | *Brevoortia tyrannus, Alosa sp* | 1101 | 8.8 |  |
| Skilletfish | *Gobiesox strumosus* | 1068 | 8.5 | Y |
| Feather blenny | *Hypsoblennius hentz* | 531 | 4.2 | Y |
| Striped bass | *Morone saxatilis* | 506 | 4.0 | Y |
| Bay anchovy | *Anchoa mitchilli* | 458 | 3.7 |  |
| Oyster Toadfish | *Opsanus tau* | 439 | 3.5 | Y |
| Seaboard goby | *Gobiosoma ginsburgi* | 419 | 3.3 | Y |
| Spot | *Leiostomus xanthurus* | 401 | 3.2 | Y |
| Black sea bass | *Centropristis striatus* | 362 | 2.9 | Y |
| Atlantic silverside | *Menidia menidia* | 299 | 2.4 |  |
| American eel | *Anguilla rostrata* | 251 | 2.0 |  |
| Scup | *Stenotomus chrysops* | 144 | 1.1 | Y |

|  |  |  |  |
| --- | --- | --- | --- |
| Atlantic herring | *Clupea harengus* | 132 | 1.1 |
| Tomcod | *Microgadus tomcod* | 108 | 0.9 |
| Bluefish | *Pomatomus saltatrix* | 104 | 0.8 |
| SUM |  | 11819 | 94.5 |

#### Phenology

There was 10-fold seasonal variation in total marine fish eDNA which roughly paralleled seasonal variation in New York Harbor water temperature (Fig 5). There was a similar seasonal pattern in daily species richness, such that no species were abundant in winter and few were common. Daily species richness for uncommon and rare species did not show a clear seasonal trend. Individual species differed in seasonality (Fig 6).

**Fig 5.**
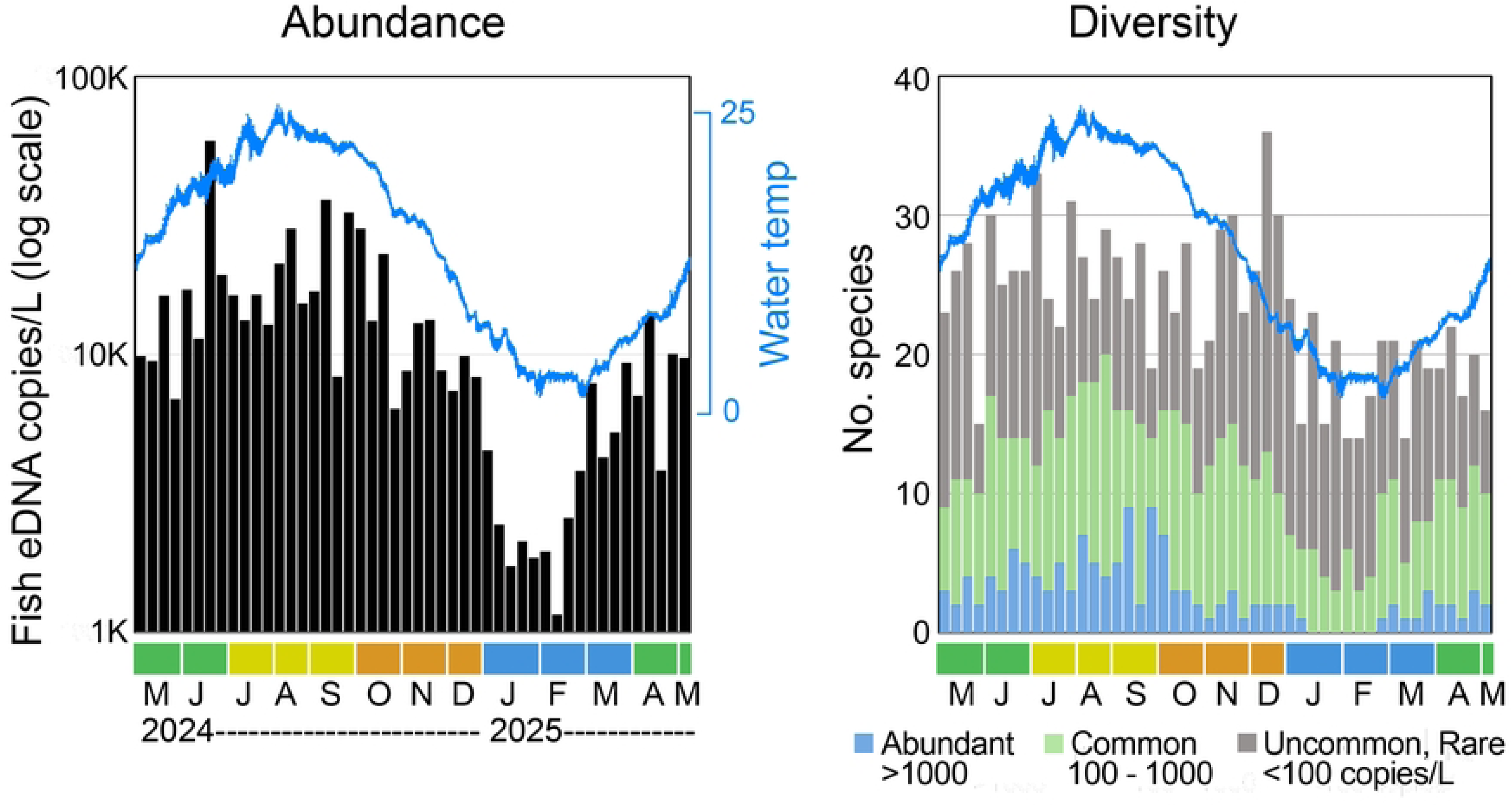
Local marine fish eDNA seasonal abundance and diversity. Each column represents one collection day. Collection months, colored by quarter, are indicated; overlay shows New York Harbor water temperature. Left, total copies/L local marine fish. Right, number of species, colorized by abundance rank on collection day as in Fig 4 (source data S3 Table).

**Fig 6.**
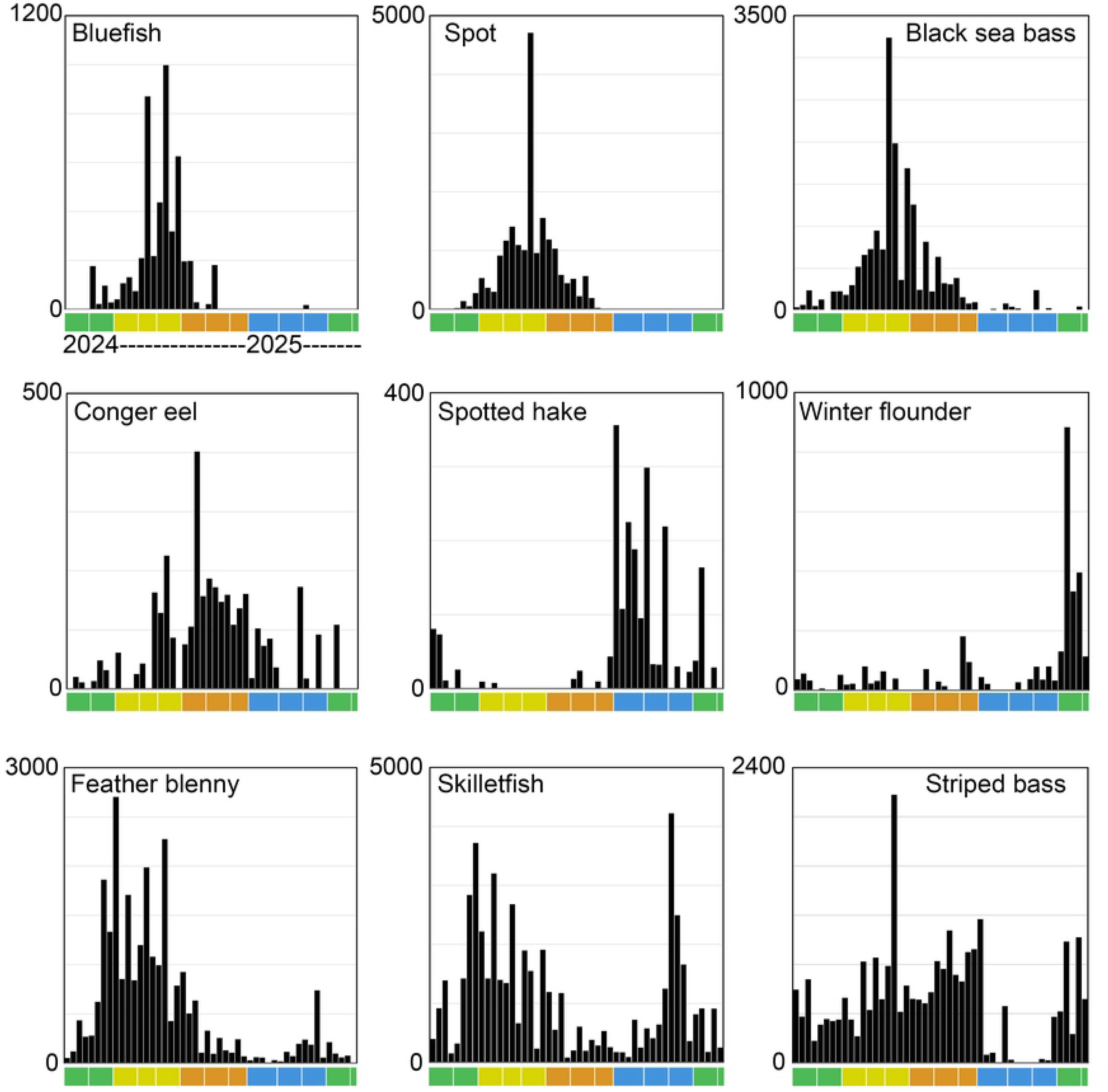
Seasonal eDNA abundance individual species. Selected species with differing phenology are shown. Each column represents one collection day; scale is copies/L. Scale differs between graphs. Color bar indicates months as in Fig 5. Scientific names, source data in S3 Table.

### Other vertebrate eDNA

Human, domesticated animal, non-fish wildlife, and nonlocal fish eDNA was commonly detected in estuary samples (Fig 2). Unlike local marine fish eDNA, daily abundances of other vertebrate eDNA categories were directly correlated to that for human eDNA, consistent with a shared wastewater source (Fig 7). Estuary eDNA correctly identified the commonest urban mammals, land birds, and household pets (Table 2) [41,42]. Dietary animal eDNA proportions closely tracked proportions in national statistics on meat and fish consumption (Fig 8) [43–46].

**Fig 7.**
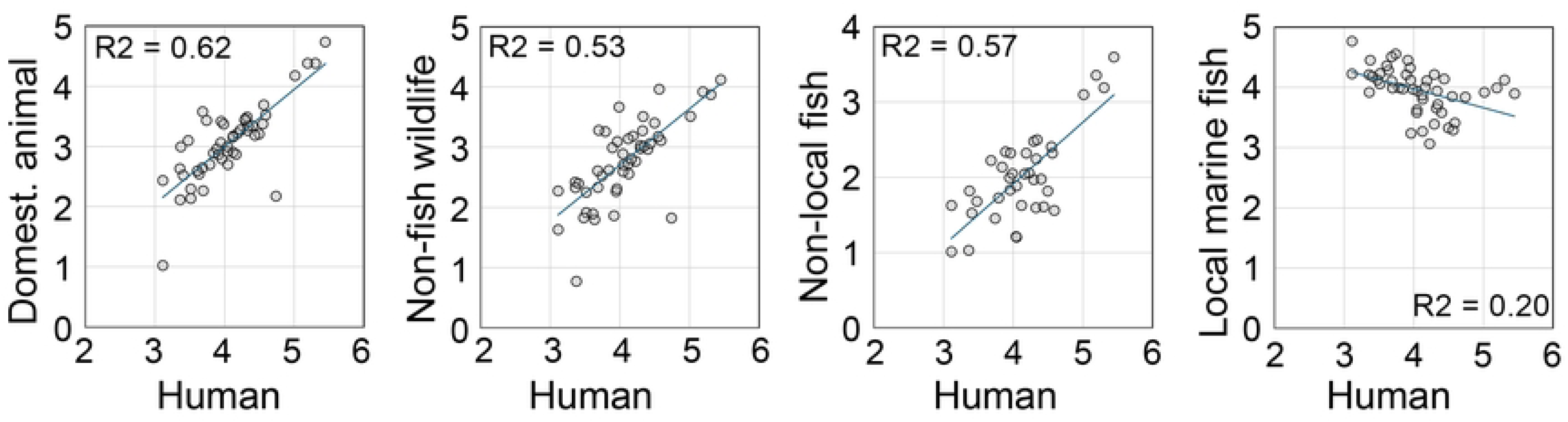
Other vertebrate eDNA abundance by category in relation to human eDNA abundance. Each point represents one collection day. Scale is log10 copies/L.

**Fig 8.**
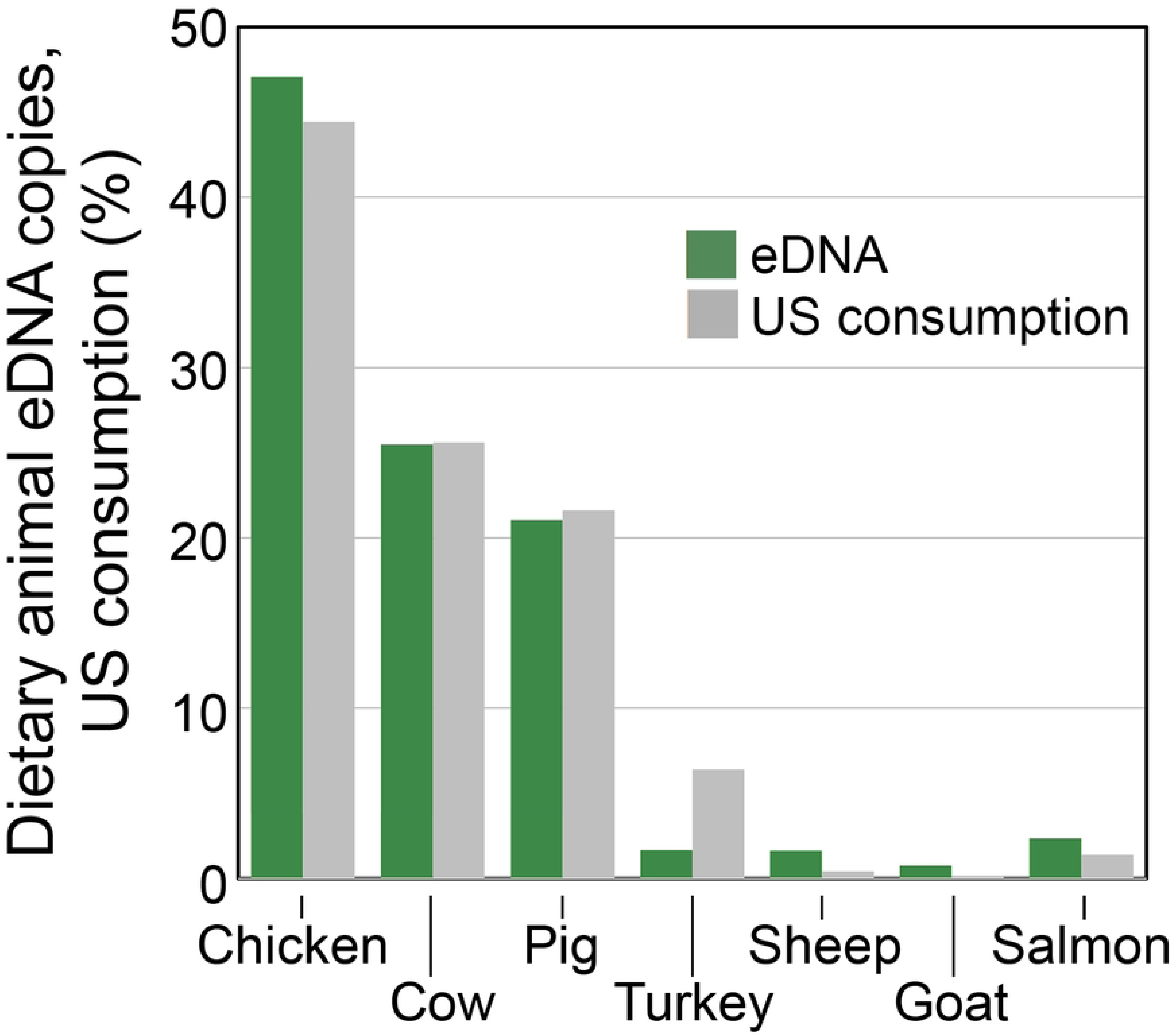
Proportions dietary animal eDNA compared with proportions US consumption per capita. Data sources in text.

**Table 2.**
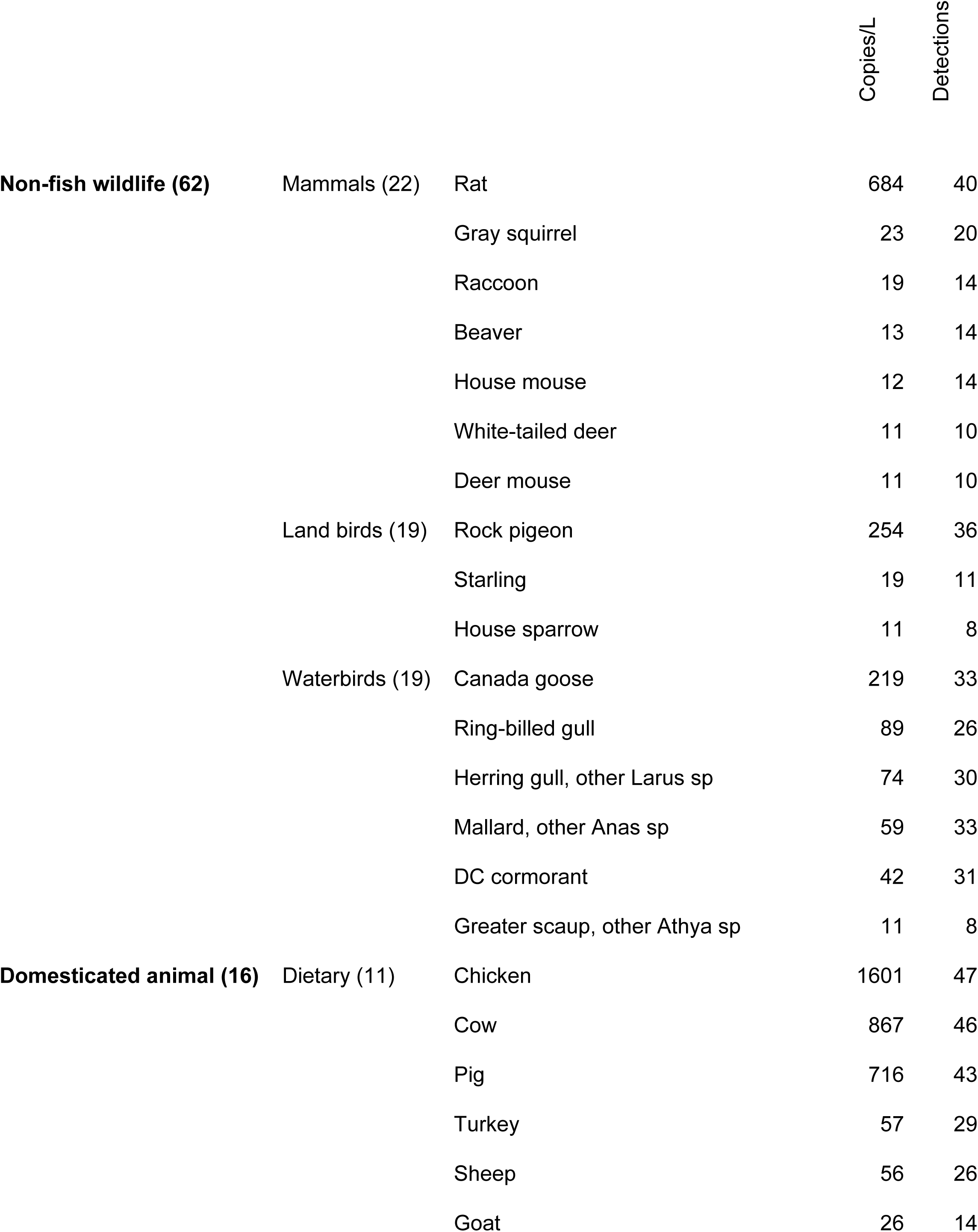
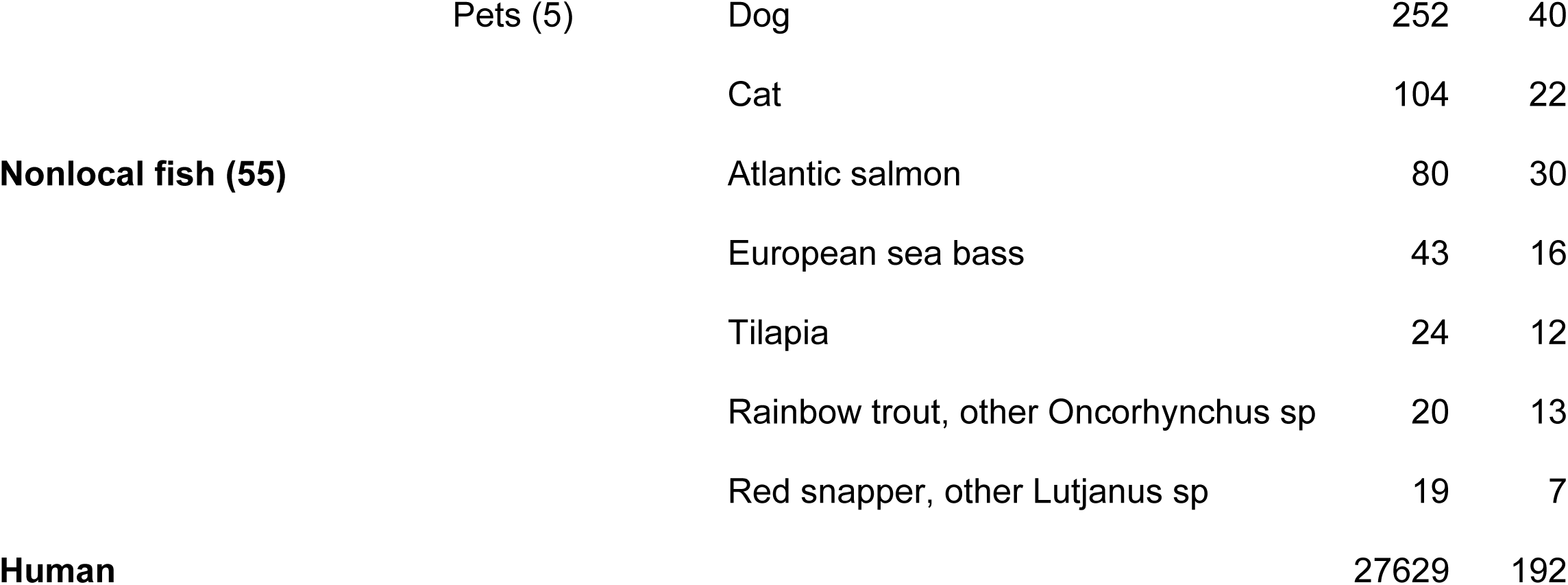
Top species other vertebrate categories.

Average copies/L and days detected (total collection days n = 48) are shown. Taxa are ranked by decreasing abundance within each category; rare species (<10 copies/L) not shown. Total detected in parentheses. Scientific names, complete dataset in S3 Table.

## Discussion

In this investigation we employed spike-in metabarcoding to quantify vertebrate eDNA weekly for one year in an urban estuary. There were two main findings. First, results supported hypothesis that eDNA indexes local marine fish abundance. eDNA followed expected fish survey characteristics including a classic hollow-curve species abundance distribution, reef specialists as top species, summer peak in total fish abundance, and species-specific seasonal patterns consistent with life histories [47–51]. Human and other wastewater eDNA was plentiful but total vertebrate eDNA levels were below the threshold expected to suppress fish assessment. Second, wastewater eDNA tracked terrestrial wildlife abundance and human consumption of meat and fish.

We propose a fish abundance scale based on order of magnitude differences in absolute eDNA concentration (Fig 2). An order of magnitude scale has been applied to rank numerical abundance of fish species [52]. Standardized numerical categories of eDNA abundance could help communicate survey findings to general as well as scientific audiences. eDNA rarity was the main limit to eDNA detection and quantification, a recognized constraint in fish eDNA surveys [53–55]. The protocol employed a PCR input of 1/20th of the DNA obtained from 1 liter of water, a similar proportion as that in other studies. Fish eDNA copies per PCR aliquot were surprisingly low (average, 625; range, 21 – 3271). This was typically sufficient to detect a dozen or so species (average, 15; range, 4 – 26). Across the one-year survey, most of 71 local marine fish species were present in fewer than 10% of PCRs (S3 Table). At the other end of the distribution, skilletfish (*G. strumosus*) and feather blenny (*H. hentz*) were newly abundant (Table 1). These taxa were rare in an eDNA survey at this site in 2016 and increased in 2022 (Fig 9) [39,56]. Both species were rare or absent in regional surveys up to 2020 [52,57–59]. Hudson River Park Fish Survey (HRPFS) corroborates recent increases [60]. Skilletfish and feather blenny were first recorded in HRPFS traps in 2020 and subsequently commonly collected. Both species favor oyster reef habitat; the current plenitude might be consequent to restoration of oyster beds in New York Harbor that began in 2015 [61].

**Fig 9.**
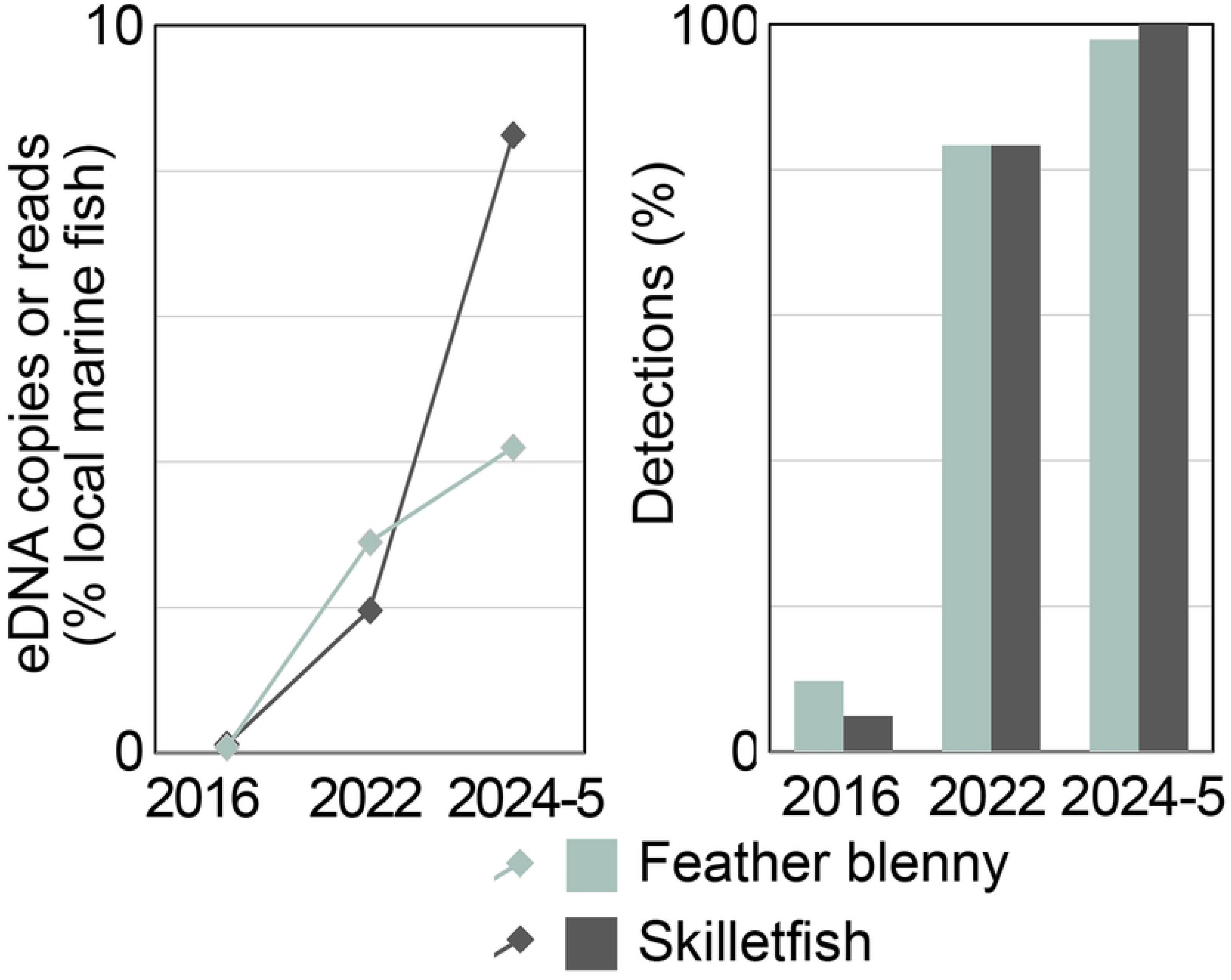
Newly abundant species. Data are from this study and previously reported East River eDNA surveys (see text for references).

Wastewater DNA offered insights. Daily levels of non-fish vertebrate eDNA correlated with daily levels of human DNA, consistent with a shared wastewater source. The highest levels of human and other vertebrate eDNA were obtained after significant rainfall, evidence that peak concentrations resulted from CSO discharge of untreated waste (S6 Fig). Whether processed wastewater contributes human and other vertebrate eDNA to the Hudson estuary is unknown. It is recently reported that dietary fish eDNA persists even after quaternary treatment of sewage [62]. Future work could examine eDNA content of untreated sewage and street runoff at CSO outfalls, at different points in the processing system, and elsewhere in the estuary. Human and domesticated animal DNA may serve as markers of human environmental impact [63].

Wastewater is often considered a nuisance in eDNA surveys because of possible suppression of fish reads by human DNA and misleading detection of dietary species [30,64]. Some studies use human blocking primers, which have unknown effects on amplification of target taxa. Present results suggest this precaution may be unnecessary, the more so if primers are selective for fish vs. other vertebrates. In this report the apparent copy number of human DNA as assayed with MiFish-U-F/R2 primers, which have multiple binding site mismatches in mammals, was about 1/1000th of that obtained with Riaz primers (S8 Table).

Rats are the most abundant wild mammal in New York City [65,66]. Monitoring rat eDNA in the estuary may help assess pest control programs. Standardizing against other terrestrial wildlife eDNA such as pigeon might compensate for the variable content of street runoff in estuary samples, or street runoff could be directly sampled. Routine estuary eDNA testing could greatly inform urban wildlife management. More than 60 non-fish wildlife species were identified including fauna rarely seen (S3 Table). Nearly all were taxa known to be resident within city limits, although transport from distant sites or local zoos cannot be excluded [41,42]. Proportions of livestock and dietary fish eDNA in wastewater closely approximated proportions in national consumption statistics. To our knowledge this is the first demonstration that wastewater eDNA reports on human diet, a finding of potential value to public health and commercial interests. The proportions of goat and sheep in East River eDNA were greater than in national data, which might reflect increased consumption by ethnic populations in New York City. This hypothesis could be explored with local sales data.

### Limitations

This report has several limitations. A concurrent gear-based survey is not available for benchmarking eDNA. Assessment of eDNA performance for fish censusing rests on features outlined above. Sampling was conducted at a single location, so findings incompletely depict lower Hudson River estuary fish populations. Nonetheless, fish eDNA diversity broadly overlapped with traditional surveys conducted in the estuary over the past 35 years [52,57–59]. Potential tidal effects were not directly assessed. Given semidiurnal tides and weekly water sample collection, consecutive samples were drawn on approximately opposite tides. Inspection did not show consistent week-to-week variation in eDNA profiles, and exploratory plots of eDNA copies/species vs. tide timetable were similarly unrevealing. The apparent absence of differences may reflect limited dispersion, as tidal excursion in the East River is less than the length of the channel [67]. Many fish species spawn in the estuary [35]. Possible contributions of spawning, larval, or juvenile fish to eDNA levels are unknown. Water temperature might alter eDNA shedding or decay rates [26]. However, such across-the-board factors insufficiently account for species-specific phenologies obtained with eDNA. eDNA persistence beyond weekly sampling intervals might blunt reporting on current fish abundance. Individual species demonstrated strong seasonal patterns consistent with known phenology, consistent with inference that the assay indexes current or recent fish density (Fig 6). In addition, peak values of human and domesticated animal eDNA associated with recent rainfall did not persist into the following week (S6 Fig). Assignment of species to categories carries uncertainties. Some local marine species are also consumed by humans—striped bass (*M. saxatilis*), for example. eRNA assays may enable distinguishing nucleic acid signals due to resident fish from those introduced by waste [68]. Nonlocal fish eDNA attributed to wastewater may have originated from extralimital strays.

Reproducibility and accuracy are desirable attributes in surveying marine biodiversity. These considerations apply to both questions raised in the Introduction—whether metabarcoding reports on eDNA levels and whether eDNA levels report on fish abundance. Replication experiments demonstrated good reproducibility in measuring eDNA levels, particularly for more abundant eDNAs. Accuracy might be distorted by differences among species in amplification efficiency. Quantitatively similar results for most species with MiFish-U-F/R2 as compared with Riaz primers suggest this is not a major factor for bony fish (Fig 3, S3 Fig). Cunner (*T. adspersus*) was an exception. The apparent concentration of cunner eDNA was about 70-fold lower with MiFish-U-F/R2 than with Riaz primers, a deficit predicted by binding site mismatch (S4 Fig). Dietary animal eDNA added evidence that the Riaz assay accurately reported relative eDNA levels.

There are limits to knowledge about abundance of marine biodiversity [69]. Censusing is inherently imprecise for fish and other nekton [70]. Given that fish SADs typically extend over four or five orders of magnitude, a reproducible protocol that indexes eDNA or fish abundance within, say, half an order of magnitude (about threefold) of the reference value would have wide use. There is no certain gold standard in benchmarking eDNA for marine fish assessment—all methods have catchability biases [71,72]. Estuary eDNA detected 22 freshwater fish species, all uncommon or rare (S3 Table); these findings may be of interest for future work. Waterbird eDNA was grouped with that of other non-fish wildlife but probably originated from birds in the estuary rather than from wastewater; this could be tested directly. The survey required about 25% effort and direct costs of about $15,000 (S9 Table). About half of effort was devoted to bioinformatics; this might be reduced by automating some procedures performed in Excel.

### Conclusion

Vertebrate eDNA metabarcoding with spike-in quantification offers a practical approach to biomonitoring in urban estuaries. This methodology holds promise as an aid to estuary fish and wildlife management and opens a window into human diet. Regular reporting of such findings would likely be of interest to both government and non-government entities [73].

Absolute quantification of eDNA levels could yield heretofore unprecedented ability to map fish abundance across diverse sites and habitats.

## Acknowledgments

We thank Jeanne Garbarino, Jen Bohn, and Jessi Hersh for generous assistance and sharing laboratory supplies and equipment, and David Thaler for helpful comments on manuscript.

## Supporting Information

**S1 Table. PCR primers, protocols.**

**S2 Table. Reference sequences.**

**S3 Table. eDNA copies per ASV by PCR, day.**

**S4 Table. eDNA copies per day by category for field samples, negative controls.**

**S5 Table. Reproducibility individual PCR and field replicates.**

**S6 Table. Reproducibility pooled PCR and field replicates.**

**S7 Table. MiFish-U-F/R2 copies/ASV.**

**S8 Table. MiFish-U-F/R2 vs Riaz copies/ASV.**

**S9 Table. Effort, costs.**

**S1 Fig. Gene block spike-in standard.**

**S2 Fig. Reproducibility of individual PCR and field replicates.**

**S3 Fig. MiFish-U-F/R2 vs Riaz pooled copies/L.**

**S4 Fig. Primer binding sites for local marine fish species detected in this study.**

**S5 Fig. Frequency of detection, copies per detection vs eDNA overall abundance.**

**S6 Fig. Human eDNA abundance and recent rainfall.**

